# Polysaccharide A-dependent opposing effects of mucosal and systemic exposures to human gut commensal *Bacteroides fragilis* in type 1 diabetes

**DOI:** 10.1101/492579

**Authors:** M. Hanief. Sofi, Benjamin M. Johnson, Radhika R. Gudi, Amy Jolly, Marie-Claude Gaudreau, Chenthamarakshan Vasu

**Affiliations:** Microbiology and Immunology, College of Medicine, Medical University of South Carolina, Charleston, SC-29425; Surgery, College of Medicine, Medical University of South Carolina, Charleston, SC-29425

**Keywords:** *Bacteroides fragilis*, gut commensal, polysaccharide A (PSA), Toll-like receptor 2, innate immunity, adaptive immunity, Type 1 diabetes, intestinal mucosa, systemic immune response, dendritic cells, T cells, regulatory T cells, gut permeability

## Abstract

*Bacteroides fragilis* (BF) is an integral component of the human colonic commensal microbiota. BF is also the most commonly isolated organism from clinical cases of intra-abdominal abscesses suggesting its potential to induce pro-inflammatory responses, upon accessing the systemic compartment. Hence, we examined the impact of mucosal and systemic exposures to BF on type 1 diabetes (T1D) incidence in non-obese diabetic (NOD) mice. The impact of intestinal exposure to BF under chemically-induced enhanced gut permeability condition, which permits microbial translocation, in T1D was also examined. While oral administration of pre-diabetic mice with heat-killed (HK) BF caused enhanced immune regulation and suppression of autoimmunity resulting in delayed hyperglycemia, mice that received HK BF by i.v. injection showed rapid disease progression. Importantly, polysaccharide-A deficient (ΔPSA) BF failed to produce these opposing effects upon oral and systemic deliveries. Further, BF induced modulation of disease progression was observed in WT, but not TLR2-deficient, NOD mice. Interestingly, oral administration of BF under enhanced gut permeability condition resulted in accelerated disease progression and rapid onset of hyperglycemia in NOD mice. Overall, these observations suggest that BF-like gut commensals can cause pro-inflammatory responses upon gaining access to systemic compartment and contribute to T1D in at-risk subjects.

## Introduction

While gut commensal microbes contribute profoundly to human health and immune system maturation^1, 2^, many recent reports also support the notion that gut commensals can contribute to the pathogenesis of autoimmune diseases like type 1 diabetes (T1D)^3-5^. It has been proposed that three mutually linked features, viz: 1) aberrant gut microbiota, 2) compromised intestinal epithelial barrier, and 3) altered immune responsiveness are linked to the contribution of commensal organisms to these diseases^6^. Gut pathobionts and pro-inflammatory immune responses can compromise gut integrity leading to the translocation of microbial components to systemic compartment, and consequently systemic inflammation and autoimmune progression^7-9^.

*Bacteroides fragilis* (BF), a Gram-negative anaerobe and an integral component of the colonic commensal microbiota of many mammals^10^, has been widely studied as a model organism to determine the influence of commensal bacteria in immune regulation^11-13^. Although BF represents a very small portion of the gut microbes^14^, it is the most commonly isolated bacterium from clinical cases of intra-abdominal abscesses^15, 16^, perhaps due to its ability to colonize the colonic crypt^11^. While capsular polysaccharides of BF are found to be essential for abscess formation^16, 17^ as well as for its growth^18^, these components, polysaccharide A (PSA) in particular, have been demonstrated to have the ability to promote immune regulation by directly acting on dendritic cells (DCs) and regulatory T cells (Tregs) in a TLR2 dependent manner^11, 19-21^. These paradoxical effects of BF suggest that intestinal mucosa and systemic compartments may produce different responses upon exposure to this commensal. Although the association between intra-abdominal abscesses and T1D incidence in humans is unknown or uninvestigated, many recent clinical and preclinical studies, including those involving large longitudinal cohorts of children of Environmental Determinants of Diabetes in the Young (TEDDY) study, associated higher prevalence of Bacteroides (phylum: Bacteroidetes) members, including BF, to T1D disease incidence / progression in humans and pre-clinical models^4, 22-32^. Further, it has been shown that increased abundance of *Bacteroides* spp precedes T1D onset in children^32^. However, if gut commensals such as BF, or other *Bacteroides* members, directly impact T1D autoimmune susceptibility has not been tested directly before.

Here, using BF as a model, we show that intestinal and systemic exposures to specific commensal organisms have opposing effects on autoimmune progression and T1D incidence in the non-obese diabetic (NOD) mouse model. While oral administration of pre-diabetic mice with heat-killed (HK) BF caused enhanced immune regulation and significant suppression of autoimmunity resulting in delayed hyperglycemia, mice that received small amounts of HK BF by i.v. injection showed rapid disease progression. Interestingly, only the wild-type (WT), but not PSA deficient (ΔPSA), BF produced opposing disease outcomes upon oral and systemic administrations. Furthermore, these effects were observed only in WT, but not TLR2 deficient, NOD mice. Most importantly, oral administration of NOD mice with BF under enhanced gut permeability recapitulated the T1D accelerating effect of systemic administration with HK BF. These results suggest that systemic access by gut symbiont such as BF, in the event of compromised gut barrier function, can act as a trigger and a catalyst for autoimmunity in T1D at-risk subjects.

## Materials and Methods

### Mice and Bacteria

Wild-type NOD/ShiLtJ (NOD), NOD-BDC2.5 TCR transgenic (NOD-BDC2.5), and NOD-*Rag 1* deficient (NOD-*Rag1*) mice were purchased from the Jackson laboratory (Maine, USA). Foxp3-GFP-knockin (Foxp3-GFP) mice in the B6 background were kindly provided by Dr. Kuchroo (Harvard Medical School). NOD-WT mice from our breeding colony were used in this study. NOD-Foxp3-GFP mice were generated by backcrossing B6-Foxp3-GFP mice to the NOD background for 12 generations. NOD-BDC2.5-Foxp3-GFP mice were generated by crossing NOD-BDC2.5 mice with NOD-Foxp3-GFP mice. NOD-TLR2 knockout (NOD-TLR2 KO) mice were provided by Dr. Chervonsky (University of Chicago). All animal studies were approved by the animal care and use committee of MUSC. To detect hyperglycemia, glucose levels in blood collected from the tail vein were determined at timely intervals using the Ascensia Micro-fill glucose test system (Bayer, USA). Mice with glucose level of >250 mg/dl for two consecutive testing were considered diabetic.

BF (ATCC 25285; NCTC 9343) and isogenic ΔPSA-BF (kindly provided by Dr. Laurie Comstock, Harvard Medical School) were cultured from single colonies in brain-heart infusion (BHI) medium under anaerobic condition for up to 72 h, diluted this initial culture 50-fold using complete medium and cultured for an additional 16 h. Bacterial cells were pelleted and washed with PBS twice by centrifugation and incubated at 65°C for 30 min for heat inactivation. Heat-killed (HK) bacterial preparations were tested for viable bacteria, if any, by plating and culturing aliquots of these preparations on BHI agar pates for 72 h. *Listeria monocytogenes* (ATCC 19111) cultured in BHI medium under aerobic condition was also processed similarly. These HK bacterial preparations were subjected to an additional quick wash in sterile distilled water to minimize salt content, and the pellet was air-dried and dry weight was determined, before suspending in PBS at 20 mg/ml concentration (w/v), as stock suspension. These stock solutions containing whole bacteria were diluted in sterile PBS as needed for in vivo studies.

### Peptide antigens, cell lines, and antibodies

Immunodominant β-cell antigen peptides [viz., 1. Insulin B (9-23), 2. GAD65 (206-220), 3. GAD65 (524-543), 4. IA-2beta (755-777), 5. IGRP (123-145), and 6. BDC2.5 TCR reactive peptide (YVRPLWVRME; referred to as BDC-peptide) were custom synthesized (Genescript Inc). Peptides 1-5 were pooled at an equal molar ratio and used as β-cell-Ag peptide cocktail as described in our earlier study^33^. PMA, ionomycin, Brefeldin A, monensin, magnetic bead based T cell and dendritic cell enrichment kits, ELISA and Magnetic bead based multiplex cytokine kits and unlabeled and fluorochrome labeled antibodies, FITC-Dextran, and other key reagents were purchased from Sigma-Aldrich, BD Biosciences, eBioscience, Invitrogen, Millipore, Miltenyi Biotec, StemCell Technologies, R&D Systems, and Biolegend. ELISA assays were read using Biorad iMark 96 well plate-reader. Luminex technology based multiplex assays were read using FlexMap3D or BioPlex 100 instruments. Real-time PCR assays were performed using transcript-specific or bacterial 16S rDNA specific primer sets and SYBR-green PCR mastermix. Flow cytometry data was acquired using FACS Calibur, FACS Verse or CyAn-ADP instruments and analyzed by using Summit or CytoBank applications.

### Treatment of mice with HK BF and peptides

Female NOD-WT mice were administered orally (500 μg/mouse/day for 15 days) or injected intravenously (i.v.; 10 μg/mouse/day every 3rd day for 15 days) with HK BF (WT or ΔPSA) and examined for blood glucose levels every week for over 30 weeks post-treatment initiation to detect hyperglycemia. Control mice were left untreated or received PBS as carrier buffer. In some experiments, mice that were treated as described above for 15 days or for 3 consecutive days, were euthanized either 24 h or 15 days post-treatment. In some experiments, NOD-WT, NOD-Foxp3-GFP, or NOD-BDC2.5-Foxp3-GFP mice were fed or i.v. injected with bacteria alone or along with β-cell antigen peptide cocktail or BDC2.5 peptide. Spleen, pancreatic LN (PnLN), Mesenteric LN (MLN), Peyer’s patch (PP), small intestinal lamina propria (SiLP) and/or large intestinal lamina propria (LiLP) cells from treated and control mice were examined for cytokine responses and/or T cell phenotypes. In some experiments, serum samples were subjected to Luminex multiplex assays to detect cytokines.

### Dendritic cells (DCs), T cells and in vitro and in vivo assays

CD11c+ DCs were enriched from spleen cells using magnetic separation kits and reagents from Miltenyi Biotec and/or StemCell Technologies. Small and large intestinal lamina propria (SiLP and LiLP) cells were prepared by collagenase digestion, density gradient centrifugation for enriching immune cells^34^, and followed by CD11c+ DCs by magnetic separation. T cells were enriched from spleens of treated and untreated control mice using negative selection kits. In antigen presentation assays, DCs that were exposed to HK-BF in vivo were pulsed with BDC2.5 peptide, washed and incubated with CD4+ T cells from NOD-BDC2.5 or NOD-BDC2.5-Foxp3-GFP mice in 96 well plates. After 4 days of culture, cells were stained for CD4 and examined for Foxp3 or GFP expression, and/or intracellular cytokines (after 4 h stimulation using PMA and ionomycin) by FACS.

### Adoptive T cell transfer experiment

Total T cells isolated from spleens of control and HK BF treated mice were transferred into 8-week old NOD-*Rag1* deficient mice (i.v.; 1×10^6^ cells/mouse) and tested for blood glucose levels every week to determine the diabetogenic properties of T cells.

### Induction of increased gut permeability

Mice were given 0.5% Dextran sulfate sodium (DSS) (w/v) in drinking water for 5 days and switched to regular water for rest of the monitoring period. Control and DSS treated mice were given HK-BF (500 μg/mouse/day) by oral gavage every day for up to 15 days starting day 0. Cohorts of these mice were euthanized on day 6, day 30, or monitored for up to 20 weeks post-treatment. To determine gut permeability, in some experiments, DSS treated and control mice were given FITC dextran (4KD; 10 mg/mouse) on day 6 by oral gavage, euthanized after 4 h, and plasma samples were examined for FITC-Dextran concentration by fluorimetry against control samples spiked with known concentration of FITC-Dextran. In some assays, whole blood collected from different groups of euthanized mice by cardiac puncture using sterile syringe into sodium citrate vials were subjected to qPCR assay using universal as well as BF specific 16S rDNA specific primer sets to determine the levels of bacterial DNA in circulation.

### Histochemical and immunofluorescence analysis of pancreatic tissues

Pancreata were fixed in 10% formaldehyde, 5-µm paraffin sections were made, and stained with hematoxylin and eosin (H&E). Stained sections were analyzed using a 0-4 grading system as described in our earlier studies^35^. At least 100 islets were examined for every group. In some experiments, pancreatic sections were stained using anti-insulin antibody followed by Alexa fluor 488- or 568-linked secondary antibodies and DAPI, and scored for insulitis based on DAPI-positive cells in islet areas and insulin expression. Insulitis was scored as described for H&E stained sections and insulin positive and negative islets were counted.

### Statistical analysis

Statistical significance (*p-value*) were calculated using GraphPad Prism and/or online statistical applications. Log-rank analysis (http://bioinf.wehi.edu.au/software/russell/logrank/) was performed to compare T1D incidence (hyperglycemia). Fisher’s exact test was used for comparing the total number of severely infiltrated islets (grades ≥3) relative to total number of islets with low or no infiltration (grades ≤2) in test vs. control groups. Mann-Whitney (nonparametric; two-tailed) or unpaired t-test (two-tailed) was employed, unless specified, for values from *ex vivo* and in vitro assays. Paired *t*-test (parametric; two-tailed) was employed for cumulative values of multiple experiments that were not performed in parallel. Values of assays done in duplicate or triplicate were averaged for each mouse or pool of mice before calculating statistical significance. A *p* value of ≤0.05 was considered statistically significant.

## Results

### Exposure of gut mucosa to BF results in suppression of autoimmune progression

Hyperglycemia is detected in female NOD mice as early as 12 weeks of age and 80-100% of the mice turn overt hyperglycemic before the age of 30 weeks. Here, we used 10-week-old pre-diabetic NOD mice to determine the impact of oral administration of HK BF on autoimmune progression and T1D incidence. As shown in Fig. 1A, oral administration of NOD mice with HK BF (500 μg/day) for 15 days resulted in delayed hyperglycemia in majority of the mice, as indicated by significantly reduced T1D incidence in treated mice compared to controls at 15 and 30 week post-treatment. While more than 80% of the mice remained diabetes-free at 15 weeks post-treatment, more than 60% of the control mice developed diabetes within this period. To determine the impact of orally administered BF on islet autoimmunity, one set of euglycemic mice were euthanized 30 days post-treatment and the pancreatic tissues were examined for degree of insulitis. Low incidence of the disease in BF treated mice was found to correlate with suppressed immune cell infiltration and profoundly higher number of islets with no or modest immune cell infiltration (grades ≤2 insulitis) (Fig. 1B).

**FIGURE 1:**
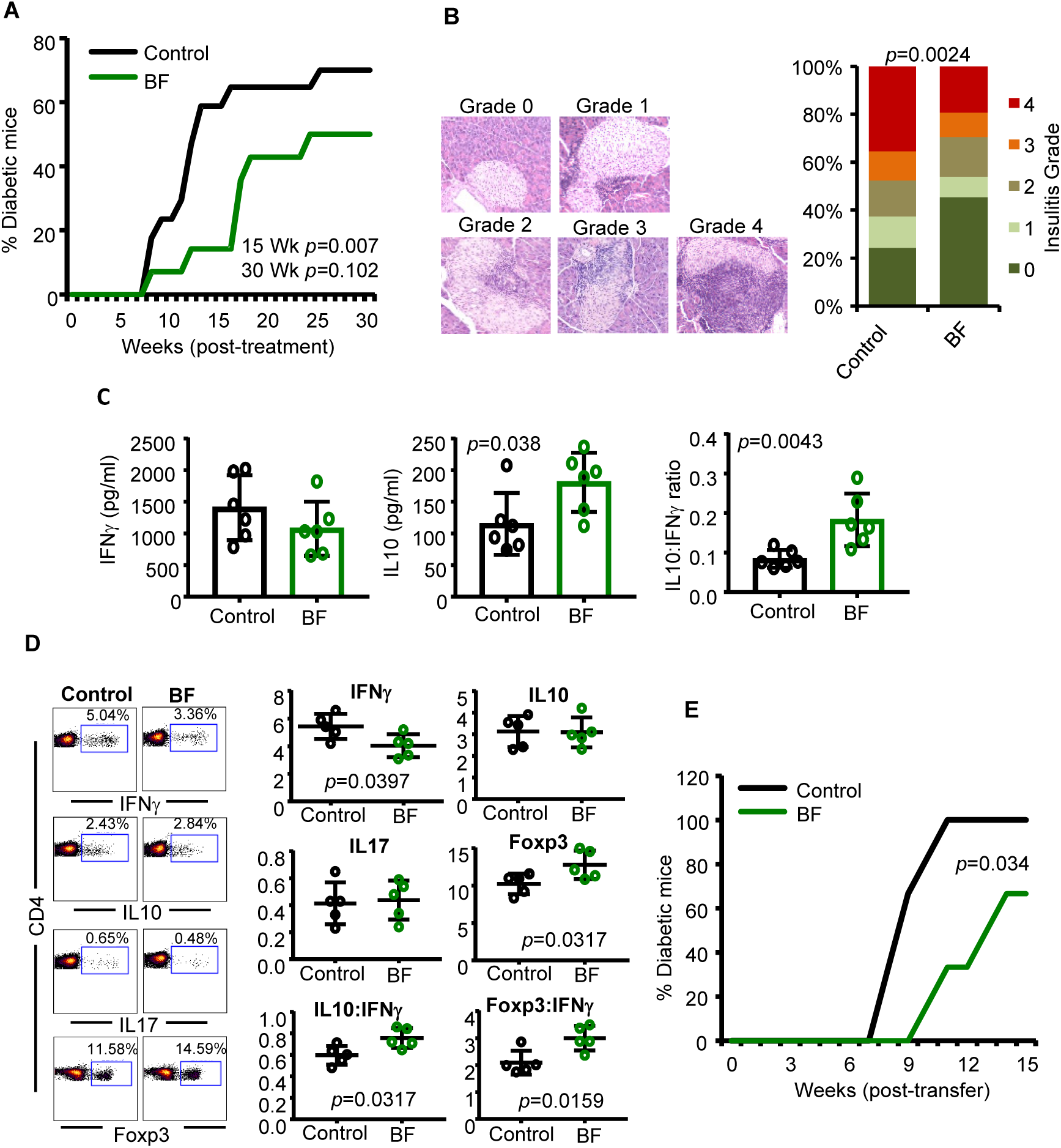
Oral administration of BF results in suppression of autoimmunity and protection of NOD mice from T1D. Ten-week-old prediabetic female NOD mice were given HK BF for 15 consecutive days (500 μg/mouse/day; dry-weight equivalent) **A)** Cohorts of control (n=17) and BF treated (n=14) mice were examined for blood glucose levels every week and the mice with glucose levels of >250mg/dl for two consecutive weeks were considered diabetic. Percentage of mice that developed diabetes at weeks post-treatment initiation are shown. Statistical significance was assessed by log-rank test for up to 15 and 30 weeks of monitoring period. **B)** Cohorts of mice (n=5/group) were euthanized 30 days post-treatment, pancreatic tissue sections were subjected to H&E staining and examined for insulitis. Examples of islets with different insulitis grades (left panel) and the percentage islets with different insulitis grades (right panel) are shown. Grades: 0 = no evidence of infiltration, 1 = peri-islet infiltration (<5%), 2= 5-25% islet infiltration, 3 = 25–50% islet infiltration, and 4 = >50% islet infiltration. At least 100 islets from intermittent sections were examined for each group. Statistical significance was assessed by Fisher’s exact test comparing relative numbers of islets with insulitis grades ≤2 and ≥3 between groups. **C)** PnLN cells of mice euthanized 15 days post-treatment (n=6 mice/group) were cultured in triplicate in the presence of β-cell Ag peptide cocktail for 48 h and the spent media was examined for self-antigen induced cytokine release by Luminex multiplex assay. Cytokine concentrations and relative ratios were compared and the statistical significance values were calculated by Mann-Whitney test. **D)** Fresh spleen cells from mice euthanized at 15 days post-treatment (n=5 mice/group) were activated ex vivo in duplicate wells with PMA and ionomycin for 4 h and stained for intracellular cytokine levels, or examined for Foxp3 levels without ex vivo activation. Percentage and relative ratios of cells that are positive for specific markers are shown and the statistical significance was assessed by Mann-Whitney test. Experiment of panels C and D were repeated using 4-6 mice/group at least once with similar statistical trends in results. **E)** Enriched splenic T cells from control and BF treated mice were adoptively transferred into 6 week old NOD-Rag1 KO mice (1×10^6^ cells/mouse) and the mice were monitored (n=6 mice/group) for hyperglycemia as described above. Log-rank test was employed for determining statistical significance.

To determine if the autoreactivity of immune cells is modulated in mice that were orally administered with HK BF, PnLN cells were activated ex vivo using β-cell antigen peptide cocktail. PnLN cells from BF treated mice produced significantly higher amounts of IL10, and showed profoundly higher IL10 to IFNγ production ratio, compared to these cells from control mice (Fig. 1C). Oral BF treatment induced modulation of systemic immune function was also assessed by examining the splenic T cells for their expression of cytokines and Foxp3. As shown in Fig. 1D, IFNγ+ T cell frequencies were lower and Foxp3+ cell frequencies were higher in BF treated mice. Further, overall IL10+:IFNγ+ and Foxp3+:IFNγ+ cell ratios were significantly higher in these mice compared to controls. To further assess if oral administration of BF impacts the diabetogenic function of systemic immune cells, splenic T cells from BF treated mice were adoptively transferred to NOD-*Rag1* mice. Fig. 1E shows that NOD-*Rag1* mice that received T cells from BF-fed mice developed hyperglycemia at a significantly slower rate compared to mice that received T cells from control mice. Overall, these observations suggest that exposure of gut mucosa to BF results in enhanced immune regulation in the systemic compartment, including pancreatic microenvironment, resulting in suppression of T1D incidence in NOD mice.

### Exposure of systemic compartment to BF results in enhanced pro-inflammatory response and accelerated T1D onset

Since BF is known to cause peritonitis, and intra-abdominal infections and abscesses upon gaining access to these compartments^15-17, 36^, and correlations of T1D autoimmunity with higher *Bacteroides* spp abundances as well as overall microbial translocation^4, 9, 22-32, 37^, we determined the effect of systemic exposure to small amounts of HK BF on autoimmune progression and T1D incidence. Ten-week-old pre-diabetic NOD mice were treated i.v. with small amounts of BF (10 μg/day) every 3 days for 15 days and monitored for hyperglycemia. As observed in Fig. 2A, mice that received i.v. injection of HK BF showed, compared to controls, rapid onset of hyperglycemia and significantly high overall diabetic incidence. Consistent with this observation, mice that were treated i.v. showed profoundly higher degree of insulitis at 30 days post-treatment (Fig. 2B). Importantly, PnLN cells from mice that received systemic treatment showed significantly high IFNγ production, and overall low IL10 to IFNγ production ratio, upon ex vivo activation using β-cell antigen peptide cocktail, compared to cells from control mice (Fig. 2C). Further, compared to control mice, significantly higher IFNγ+ and lower Foxp3+ T cell frequencies were detected in the spleen of mice that were exposed to BF systemically (Fig. 2D). In addition, *NOD-Rag1* mice that received splenic T cells from BF injected mice showed relatively faster onset of hyperglycemia, albeit not significant statistically, compared to those that received cells from control mice (Fig. 2E).

**FIGURE 2:**
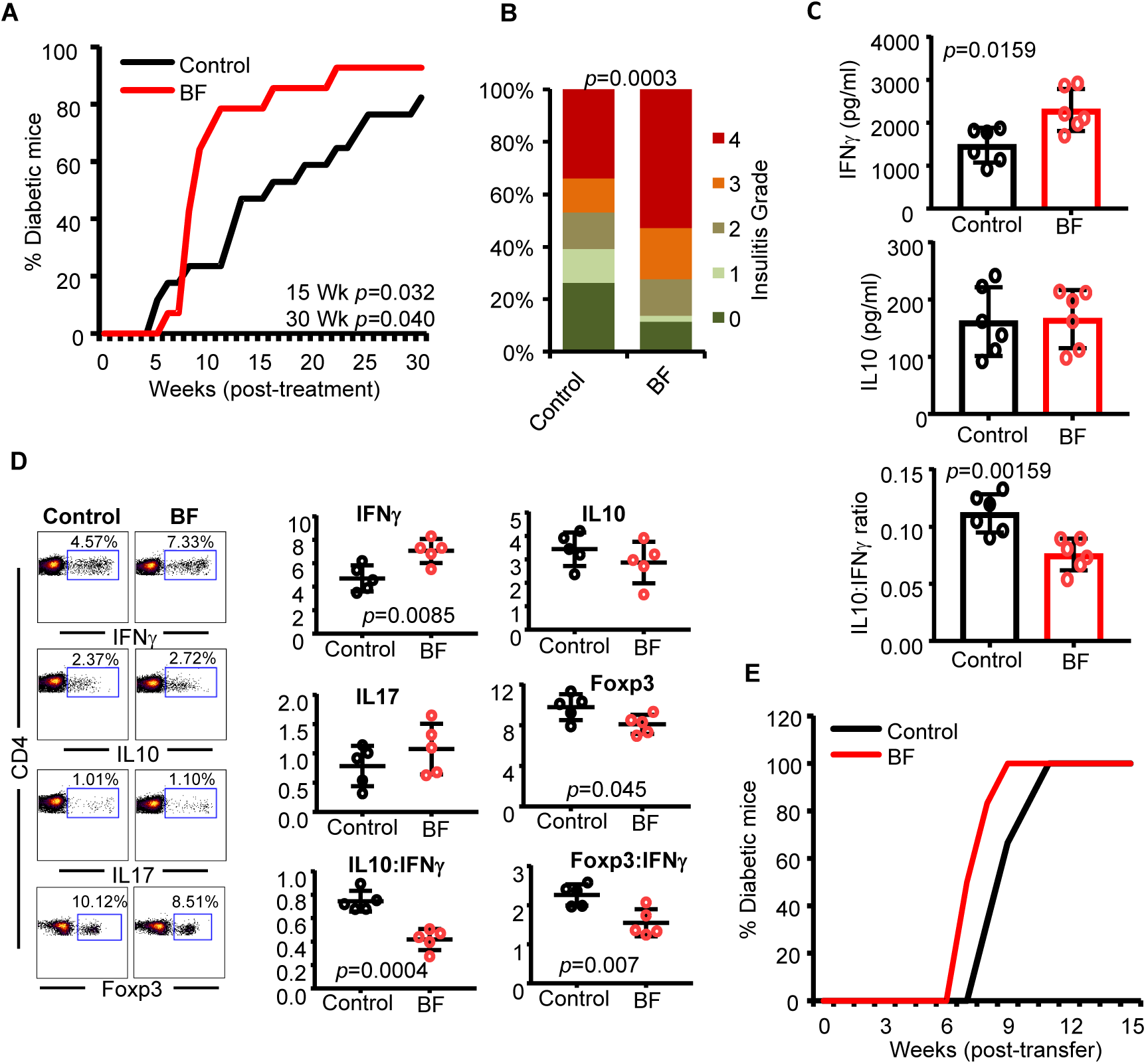
Systemic (i.v.) administration of BF results in aggravated autoimmune response and rapid T1D onset in NOD mice. Ten-week-old pre-diabetic female NOD mice were given heat-killed BF every 3rd day during 15 day period (10 μg/mouse/injection; dry-weight equivalent) by i.v. injection and studied as detailed for Fig. 1. **A)** Cohorts of control (n=17) and BF treated (n=14) mice were monitored for hyperglycemia as described for Fig. 1. Statistical significance was calculated by log-rank test for up to 15 and 30 weeks of monitoring periods. **B)** Cohorts of mice (n= 5/group) were euthanized 30 days post-treatment and pancreatic tissue sections were examined for insulitis, and the statistical significance was calculated by Fisher’s exact test. **C)** PnLN cells of mice (n=6/group) were examined for self-antigen induced cytokine levels by Luminex multiplex assay and the statistical significance values were calculated by Mann-Whitney test. **D)** Fresh spleen cells from BF treated mice (n=5/group) and controls, 15 days post-treatment were examined for intracellular cytokines and Foxp3 levels, and the statistical significance was calculated by Mann-Whitney test. Experiment of panels C and D were repeated using 4-6 mice/group at least once with similar statistical trends in results. **E)** Enriched splenic T cells from control and BF treated mice were adoptively transferred into 6 week old NOD-Rag1 KO mice (n=6/group) and monitored for hyperglycemia.

The systemic effects of i.v. injection with BF in NOD mice (Fig. 2) are opposite to that produced by oral treatment with this agent (Fig. 1). To determine if such contrasting effects are produced by other gut bacteria, pre-diabetic NOD mice were treated with HK *Listeria monocytogenes* (LM) by oral gavage or i.v. injection and examined for T1D incidence and insulitis. As observed in Fig. S1, orally and systemically administered HK LM produced disease incidence and insulitis trends in NOD mice similar to that of HK BF treated mice, however these observations were, unlike that from BF treated mice, not significant statistically by log-rank test and Fisher’s exact test.

### Orally and systemically administered BF induces distinct immune responses in the gut mucosa and spleen

Since NOD mice that were treated with HK BF orally and systemically showed opposing effects on insulitis and hyperglycemia onset, intestinal and splenic immune responses of mice that were exposed to BF for 3 days were determined. qPCR assay of distal ileum and distal colon from mice that received BF orally and spleen cells from mice that received BF systemically show that BF exposure triggers the expression of mRNA for a large number of cytokines, chemokines, and immune regulatory enzyme such as Aldh1A2 in both systemic and intestinal compartments. Notably, compared to controls, higher *Il10* mRNA expression was induced in BF exposed intestine as well as spleen. However, most intriguingly, *Il6* expression was profoundly higher in the spleen of mice that received i.v. injection, but significantly low in the intestine of oral treatment recipients, of BF compared to their control counterparts. Further, BF exposure associated *Il12* and *Il1b* mRNA expressions were significantly higher in the spleen of i.v. injected mice, but not in the intestine of orally fed mice (Fig. 3A).

**FIGURE 3:**
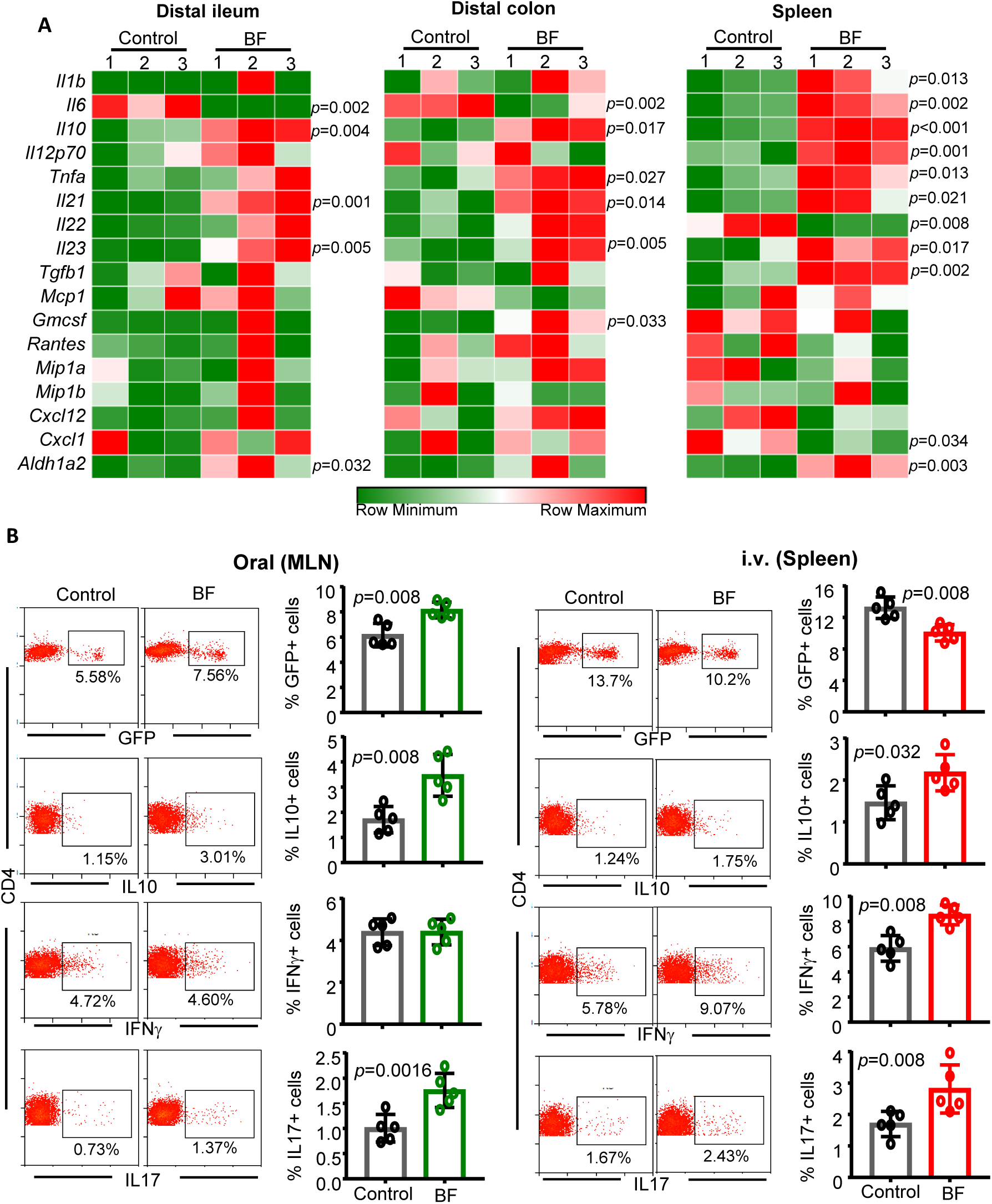
Orally and systemically administered BF induces distinct immune responses. Eight week old female NOD mice were given HK BF for 3 consecutive days by oral gavage (500 μg/mouse/day) or by i.v. injection (10 μg/mouse/injection). **A)** On day 4, one set of mice were euthanized and cDNA prepared from distal ileum and distal colon of control and BF fed (oral) mice and spleen cells from control and BF injected (systemic) mice were subjected to qPCR assay and the expression levels of cytokines and non-cytokine factors were compared. Expression levels relative to β-actin expression were plotted as heatmaps with Morpheus application. n= 3 mice/group and the assay was performed in triplicate and for each mouse the average values were used for each lane. This experiment was repeated using 3 mice/group at least once with similar statistical trends in outcomes. **B)** A set of NOD-Foxp3-GFP mice from a similar experiment (treated with BF for 3 days) were euthanized on day 7. MLN cells of BF fed and spleen cells from BF injected and control mice were examined for GFP+CD4+ cells, or ex vivo stimulated with PMA and ionomycin for 4 h and examined for intracellular cytokines, by FACS. Representative FACS plots (left panels) and plots showing percentage and relative ratios of cells that are positive for specific markers (right panels) are shown. n= 5 mice/group and the assay was performed in duplicate for each mouse. This experiment was repeated at least twice (using 3 or 4 mice/group) with similar statistical trends in outcomes. The statistical significance was assessed by Mann-Whitney test for all panels. **Notes:** Fig. S2 shows Foxp3, IFNγ, IL10, IL17 positive CD4 T cell frequencies from spleen of orally treated and MLN of i.v. treated mice used for panel B. Fig. S3 shows Foxp3, IFNγ and IL10 positive CD4+ T cell frequencies of PP, SiLP, LiLP and spleen of mice that received BF orally and spleen and PP of mice that received i.v. injection of BF for an extended period.

To determine the impact of BF induced intestinal and systemic immune responses on T cell phenotype, mesenteric lymph nodes (MLN) from NOD-Foxp3-GFP mice that received oral administration and spleens from mice that received systemic administration of BF for 3 days were compared for Foxp3 (GFP) and intracellular cytokine positive T cell frequencies on day 7. As observed in Fig. 3B, while MLN of mice that received BF orally showed significantly higher frequencies of GFP+, IL10+ and IL17+ T cells compared to their control counterparts, spleens of mice that received systemic BF showed profoundly lower frequencies of GFP+ cells, and higher frequencies of IFNγ+ cells compared to controls. Further, mice that received BF systemically for 3 days also showed higher splenic IL17+ and IL10+ cell frequencies. Spleen cells and MLN cells from mice that were treated for 3 days with HK BF orally and systemically respectively showed only a modest effect on T cell phenotype (Fig. S2). In a separate experiment, pre-diabetic mice were also subjected to prolonged (15 day) oral and systemic treatments as described for Figs. 1 and 2, euthanized within 24 h, and examined for T cell phenotype. Fig. S3 shows significantly higher Foxp3+ and IL10+, and lower IFNγ+ CD4+ cells, in the gut associated lymphoid tissues (GALT) as well as spleen of mice that received prolonged treatment. On the other hand, prolonged systemic treatment caused significantly reduced Foxp3+ and IL10+, and increased IFNγ+, CD4+ T cell frequencies primarily in the spleen, but not in the PP. Overall, these observations, along with the cytokine expression profile data of Fig. 3A, suggest an overall regulatory immune response by gut mucosa and pro-inflammatory immune response by systemic immune cells upon exposures those compartments to BF.

### BF exposed intestinal and splenic APCs produce opposing effects on Tregs upon antigen presentation ex vivo and in vivo

Since exposure to HK BF resulted in contrasting immune profiles in the intestinal mucosa and systemic compartment, impacts of exposure to BF on intestinal and splenic antigen presenting cells and antigen specific T cell activation were determined. NOD mice were treated HK BF for 3 consecutive days and euthanized on day 4 as described for Fig. 3A. CD11c+ dendritic cells (DC) were enriched from the small intestine of BF fed and spleen of BF injected mice along with respective control mice, and used in qPCR and antigen presentation assays. Although low overall levels, relative to abundantly expressed actin, perhaps due to weak activation of a small fraction of total DC preparation in vivo, fig. 4A shows cells isolated from the intestine of mice that received BF orally expressed significantly less *Il6* and *Il12*, and more *Il10* and *Tnfa* mRNA compared to cells from control mouse intestines. On the other hand, splenic DCs of mice that received systemic treatment with BF expressed higher *Il6, Il12, Il10* and *Tnfa* mRNA compared to their counterparts from control mice. However, splenic DCs from orally treated and intestinal DCs from i.v. treated mice showed comparable cytokine expression profiles compared to that of their control counterparts (Fig. S4).

**FIGURE 4:**
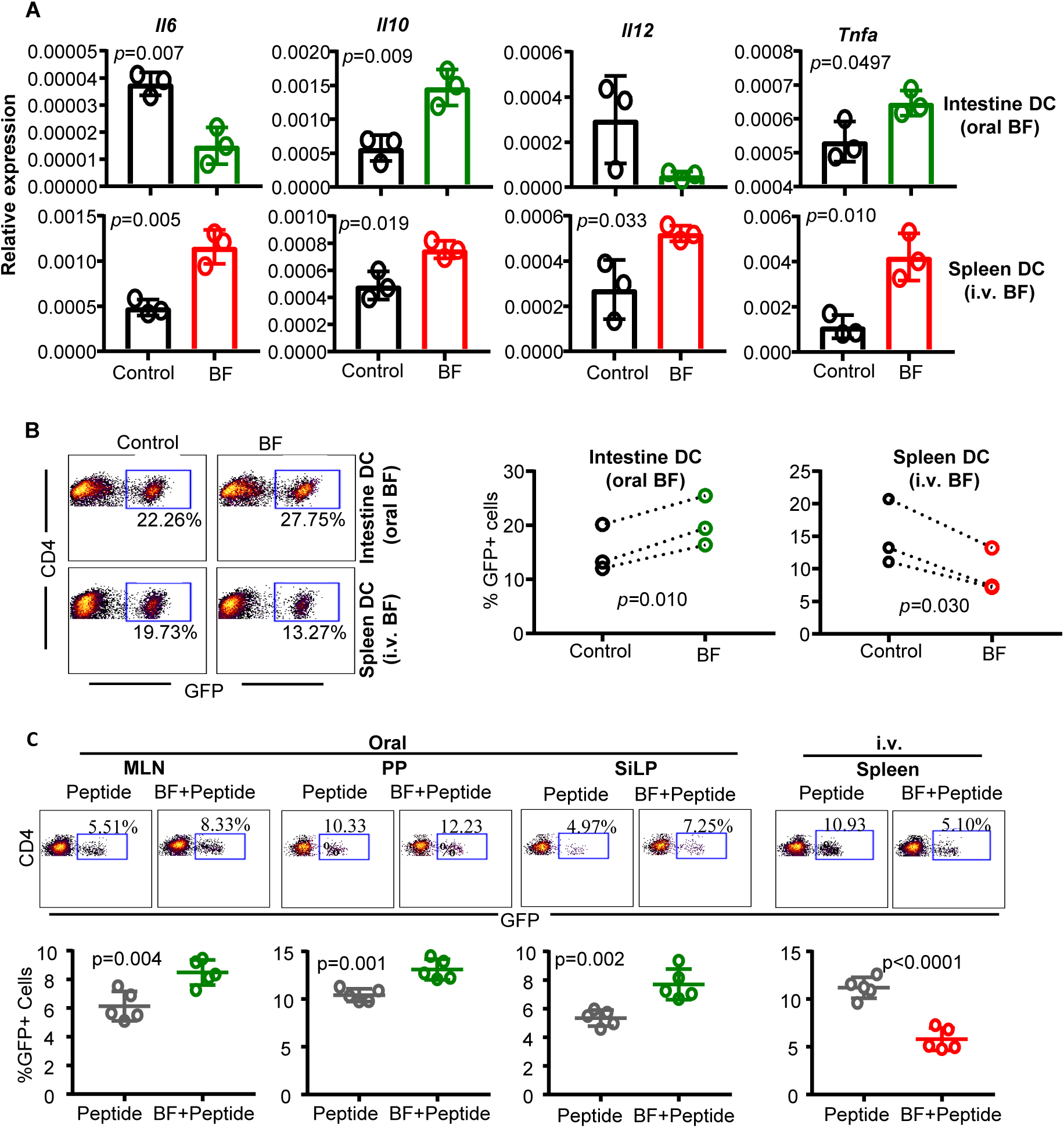
BF exposed intestinal and splenic APCs produce opposing effects on Tregs upon antigen presentation ex vivo and in vivo. Eight week old female NOD mice were given HK BF for 3 consecutive days by oral gavage or by i.v. injection as described for Fig. 3. **A)** One set of mice were euthanized within 24 h of treatment and CD11c+ DCs were enriched from the ileum of control and BF fed (oral) mice and spleen cells of control and BF injected (systemic) mice and subjected to qPCR assay to detect the expression levels of cytokines. n= 3 parallel independent assays using DCs enriched and pooled from 2 mice/group, and each assay was performed in triplicate. The statistical significance was assessed by t-test (unpaired; parametric; two-tailed). This experiment was repeated at least once with similar statistical trends in outcomes. **Notes:** Fig. S4 shows expression levels of various factors in DCs isolated from the intestine of i.v. treated and spleen of orally treated mice used for panel A. **B)** DCs were enriched from cohorts of mice similar to that of panel A, pulsed with BDC2.5 peptide and cultured with T cells enriched from BDC2.5-Foxp3-GFP mice for 4 days and examined for GFP+CD4+ cells by FACS. n= 3 independent assays using DCs enriched and pooled from at least 2 mice/group in each. Each assay was performed in triplicate and the cumulative values of three independent assays are shown. The statistical significance value was determined by t-test (paired; parametric; two-tailed) which compared control and BF groups within each assay as indicated by the dotted lines. DCs isolated from large intestine showed similar properties in terms of cytokine expression and T cell activation (not shown). **Notes:** Fig. S5A shows GFP+CD4+ T cell frequencies in cultures where T cells were activated using intestinal DCs from BF injected and spleen DCs from BF fed mice. Fig. S5B shows cytokine levels in the culture supernatants from the assays of Fig. 4B, tested by ELISA. **C)** BDC2.5-Foxp3-GFP mice that were treated for 3 days as described above received BDC2.5 peptide by oral gavage (20 μg/mouse) or by i.v. injection (2 μg/mouse) on days 2 and 3. These mice were euthanized on day 7, GFP+CD4+ T cell frequencies in MLN, PP and SiLP of orally treated and spleen cells from i.v. treated mice were determined by FACS. n= 5 mice/group and the assay was performed in duplicate for each mouse. This experiment was repeated at least once (using 4 mice/group) with similar statistical trends in outcomes. The statistical significance was assessed by t-test (unpaired; parametric; two-tailed).

To examine the impact of ex vivo antigen presentation by APCs that were exposed to BF in vivo, BDC2.5 peptide pulsed DCs were cultured with T cells from NOD-BDC2.5-Foxp3-GFP mice and the Foxp3 (GFP)+ T cell frequencies were determined in these cultures. As shown in Fig. 4B, while intestinal DCs from mice that received BF orally caused a significant increase in GFP+ CD4 T cells compared to these cells from the control mice, GFP+ CD4 T cell frequencies were significantly lower in cultures where splenic DCs from BF injected mice were used. In addition, as shown in Fig. S3, activation of antigen specific T cells by intestinal DCs from BF fed mice resulted in relatively lower IFNγ and significantly high IL10 release, and an overall higher IL10:IFNγ ratio, in the cultures. On the other hand, activation of T cells by splenic DCs from BF injected mice resulted in higher IFNγ and lower IL10 release, and profoundly low IL10 to IFNγ ratio, in the cultures. Of note, effects of splenic DCs from BF fed mice and intestinal DCs from BF injected mice on T cells were comparable to that of respective DC preparations from control mice (Fig. S5A). Overall, these observations along with the data of Figs. 1, 2, S2, S3 and S5, suggest that while short-term systemic and oral administrations with HK BF have significant impacts primarily on systemic lymphoid tissues and GALT respectively, immune modulation associated with prolonged exposure of gut mucosa to BF is also detectable in systemic organs including PnLN and spleen.

To determine the impact of antigen presentation by BF exposed APCs in vivo, NOD-BDC2.5-Foxp3-GFP mice were treated with BDC2.5 peptide alone or along with BF orally or systemically and examined for GFP+ CD4 T cell frequencies in GALT and spleen respectively. Compared to the BDC2.5 peptide alone recipients, mice that received oral gavage of BDC2.5 peptide along with BF showed significantly higher frequencies of GFP+ CD4 cells in the GALT including MLN, PP and SiLP (Fig. 4C). However, mice that received systemic injection of BDC2.5 peptide along with BF had profoundly diminished frequencies of GFP+ CD4 T cells in the spleen than those that received BDC2.5 peptide alone. These observations, in association with the results on T1D incidences in BF treated mice (shown in Figs. 1 & 2), indicate that while BF-induced immune response in the gut mucosa promotes immune regulation, systemic immune response against BF promotes acceleration of the autoimmune process in T1D.

### Oral or systemic administration of PSA deficient (ΔPSA)-BF failed to impact autoimmune progression and T1D incidence in NOD mice

Since PSA has been considered as the major symbiotic factor and host immune modulator of BF^11^, we determined if isogenic ΔPSA mutant strain^38^ of BF produces immune responses in the intestinal and systemic compartments similar to that by BF. Cohorts of mice were treated orally or systemically with HK BF and ΔPSA-BF for 3 days and cytokine expression profiles in the intestine and spleen respectively were determined by qPCR. Fig. S6 shows that mice that received ΔPSA-BF orally for 3 days, compared to BF recipients, showed significantly higher expression of *Il6* and *Il12* mRNA, and lower expression of *Il10* mRNA in the intestine. On the other hand, spleen cells of mice that received ΔPSA-BF systemically showed diminished expression of mRNA for most cytokines, albeit not statistically significant except for *Il10*, and profoundly increased expression of *Il23* compared to spleen cells from BF recipient counterparts.

To determine the intestinal and systemic impacts of PSA deficiency in BF on T1D incidence, pre-diabetic female NOD mice were given HK BF or its ΔPSA mutant by oral gavage or by i.v. injection for 15 days as described for Figs. 1 and 2. Pancreatic tissues from cohorts of these mice euthanized 30 days post-treatment were examined for the degree of immune cell infiltration and insulin positive islets. Fig. 5A shows, as observed for Fig. 1B, significant suppression of insulitis as well as higher frequencies of insulin positive islets in mice that were orally administered with BF. Mice that received BF systemically showed, as observed for Fig. 2B, more severe insulitis as well as lower frequencies of insulin positive islets. Interestingly, mice that received ΔPSA orally and systemically showed islet function and immune cell infiltration comparable to that of control mice. Examination of PnLN cells showed that Foxp3+ T cell frequencies were significantly increased upon oral treatment and diminished upon systemic administration compared to controls only in BF, but not ΔPSA-BF, recipient mice (Fig. 5B). Further, only PnLN cells from BF, but not ΔPSA-BF, recipient mice showed significant differences in the β-cell antigen activation induced release of IL10 and IFNγ as compared to these cells from control mice (Fig. 5C). Moreover, systemic and intestinal exposures to ΔPSA-BF induce different cytokine expression profiles compared to that induced by WT BF (Fig. S6). In concurrence with these observations on immune cell phenotypes, insulitis and insulin positive islets, the timing of hyperglycemia and the disease incidence rate in NOD mice that received ΔPSA-BF orally and systemically were comparable to that of their control counterparts (Fig. 5D). Overall, these studies using HK ΔPSA-BF suggest that PSA is the primary factor responsible for the direct effect of BF induced opposing responses of gut mucosa and systemic immune cells, and the overall disease outcomes.

**FIGURE 5:**
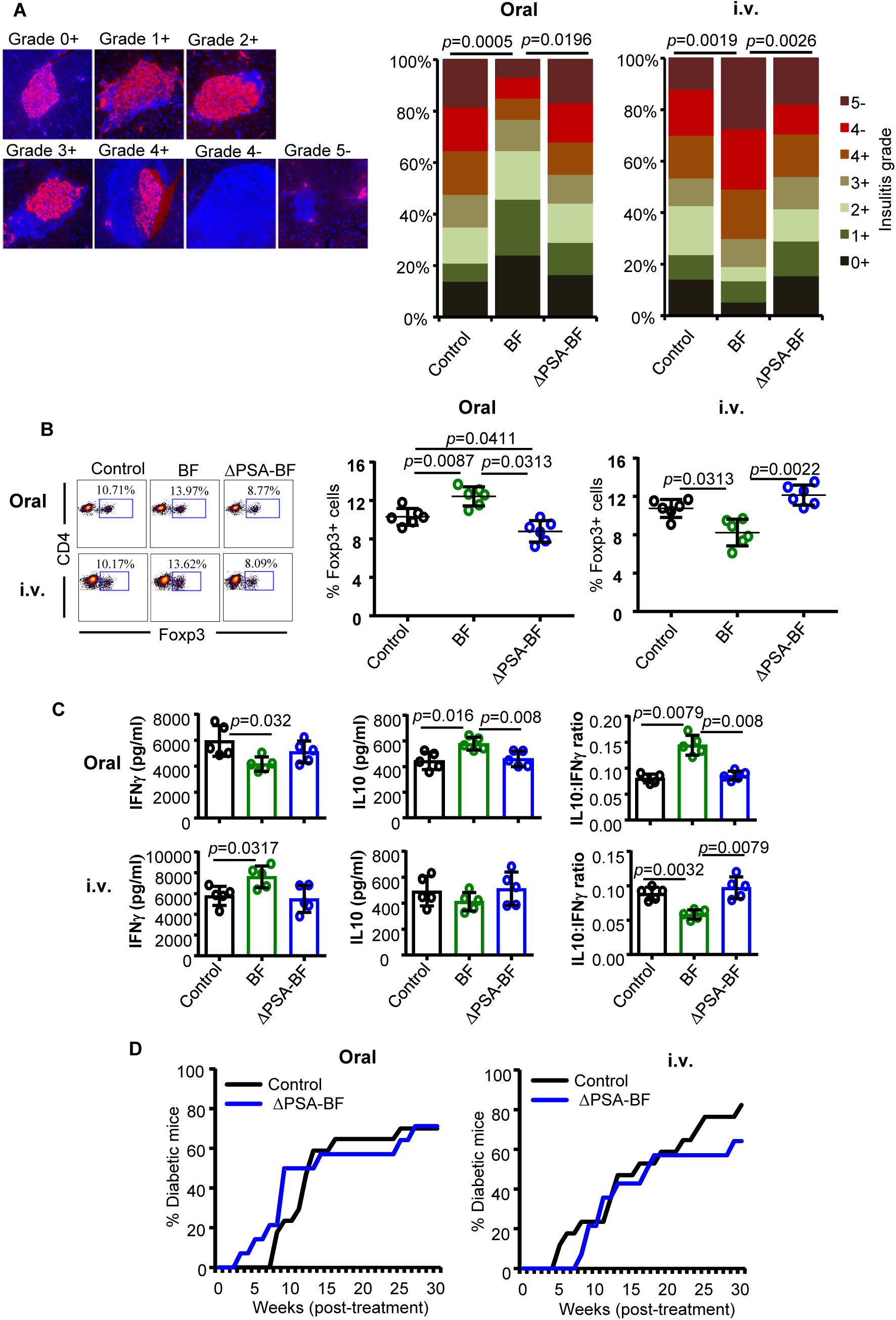
Oral or systemic administration of ΔPSA-BF has no major impact on autoimmune progression and T1D incidence in NOD mice. Ten-week-old female NOD mice were given heat-killed BF or ΔPSA-BF by oral gavage or by i.v. injection and studied as described for Figs. 1&2. **A)** Cohorts of (4 mice/group) were euthanized 30 days post-treatment, pancreatic tissue sections were stained using anti-insulin Ab (red) and the nuclear stain DAPI (blue) and examined for insulitis and insulin positive islets. Insulitis was graded 0-5 based on islet structure and immune cell infiltration (based on DAPI) and insulin staining (+ or -). 0+ (no infiltration/insulin+), 1+ (<5% infiltration/insulin+), 2+ (5-25% infiltration/insulin+), 3+ (25- 50% infiltration/insulin+), 4+ (50-100% infiltration/insulin+), 4- (50-100% infiltration/insulin-), and 5- (only islet remnants left/insulin-). At least 100 islets from multiple intermittent sections were examined for each group. Statistical significance was calculated by Fisher’s exact test comparing relative numbers of islets with insulitis grades ≤2+ and ≥3+ between control vs BF and BF vs ΔPSA-BF groups. **B)** PnLN cells of mice (n=5/group) euthanized 15 days post-treatment were examined for Foxp3+CD4+ T cell frequencies by FACS, and the statistical significance was calculated by Mann-Whitney test. **C)** PnLN cells (n=5 mice/group) were also cultured overnight in the presence of soluble anti-CD3 antibody for 24 h and the supernatants were tested for cytokine levels by Luminex multiplex assay. Cytokine levels and relative ratios are shown and the statistical significance was calculated by Mann-Whitney test. Experiment of panels C and D were repeated at least twice (using 3 or 4 mice/group) with similar statistical trends in results. **D)** Cohorts of control (n=17) and BF treated (n=14) mice were monitored for hyperglycemia as described for Figs. 1 and 2. Statistical significance was calculated by log-rank test. **Notes:** This set of experiments was carried out in parallel to the experiments of Figs. 1A and 2A and the BF group was not included in Fig 5D to avoid repeated presentation of the same data. Fig. S6 shows the cytokine mRNA expression profile of intestinal tissues from mice that were fed or injected with BF and ΔPSA-BF for 3 days.

### TLR2 deficient NOD mice, unlike WT mice, failed to show opposing T1D modulatory effects upon systemic and oral administration of BF

Interaction of BF-PSA with TLR2 plays a key role in the symbiotic function of BF^11^. Therefore, the contribution of TLR2 in protecting NOD mice from T1D upon oral administration and accelerating the disease progression upon systemic administration of HK BF was determined using NOD-TLR2-KO mice. Ten week old TLR2 KO and WT NOD mice were given HK BF orally or i.v. as described for Figs. 1 and 2 and monitored for hyperglycemia. Fig. S7 shows that, as reported by others^3, 5, 39^, NOD-TLR2 KO mice developed hyperglycemia at a relatively lower pace than the WT mice. However, unlike WT NOD mice, neither systemic nor oral treatment with BF produced a modulatory effect on the onset of hyperglycemia or T1D incidence in TLR2 KO mice. Similarly, examination of pancreatic tissues of similarly treated mice 30 days post-treatment showed that control and BF treated NOD-TLR2KO mice had comparable insulitis. These results, in conjunction with the data of Figure 5, suggest a direct mechanistic role for PSA-TLR2 interaction in the opposing T1D modulatory effects observed upon exposure of gut mucosa and systemic compartment to BF.

### Oral administration of HK BF under enhanced gut permeability results in accelerated T1D

Compromised gut epithelial barrier function and translocation of microbial components to the systemic compartment can occur under many circumstances including stress and medications. Recent reports have suggested that leaky-gut and microbial translocation may contribute to initiation and/or progression of different autoimmune conditions^8, 9, 37, 40-43^. Since systemic administrations of HK BF caused rapid disease onset in NOD mice, we determined the impact of oral administration of this agent under enhanced gut permeability conditions. To enhance gut permeability, we used a chemical approach of epithelial barrier injury. Pre-diabetic NOD mice were given low-dose Dextran sulfate sodium (DSS) in drinking water before oral administration of HK BF. DSS treated mice showed higher translocation of orally administered FITC-dextran as well as microbial DNA in to blood (Fig. S8) suggesting that they have higher gut permeability. While HK BF recipients, as observed in Fig.1, showed protection from T1D, DSS treatment alone did not impact the disease progression in NOD mice of our SPF facility (Fig. 6A). However, compared to mice that received DSS alone, mice that received both DSS and HK BF showed rapid progression of the disease and early onset of hyperglycemia. Further, pancreas of mice that received both DSS and BF showed significantly higher number of islets with severe insultis compared to DSS recipient controls (Fig. 6B). Examination of PnLN cells revealed significantly low Foxp3+ T cell frequency (Fig. 6C) and profoundly higher β-cell antigen induced IFNγ production (Fig. 6D) in DSS and BF treated mice compared to DSS treated controls. Overall, these results suggest that while transient translocation of gut microbial components normally may not have a profound impact on autoimmune progression, systemic access to specific gut microbes such as BF, especially under gut permeability compromised conditions, can accelerate the disease process.

**FIGURE 6:**
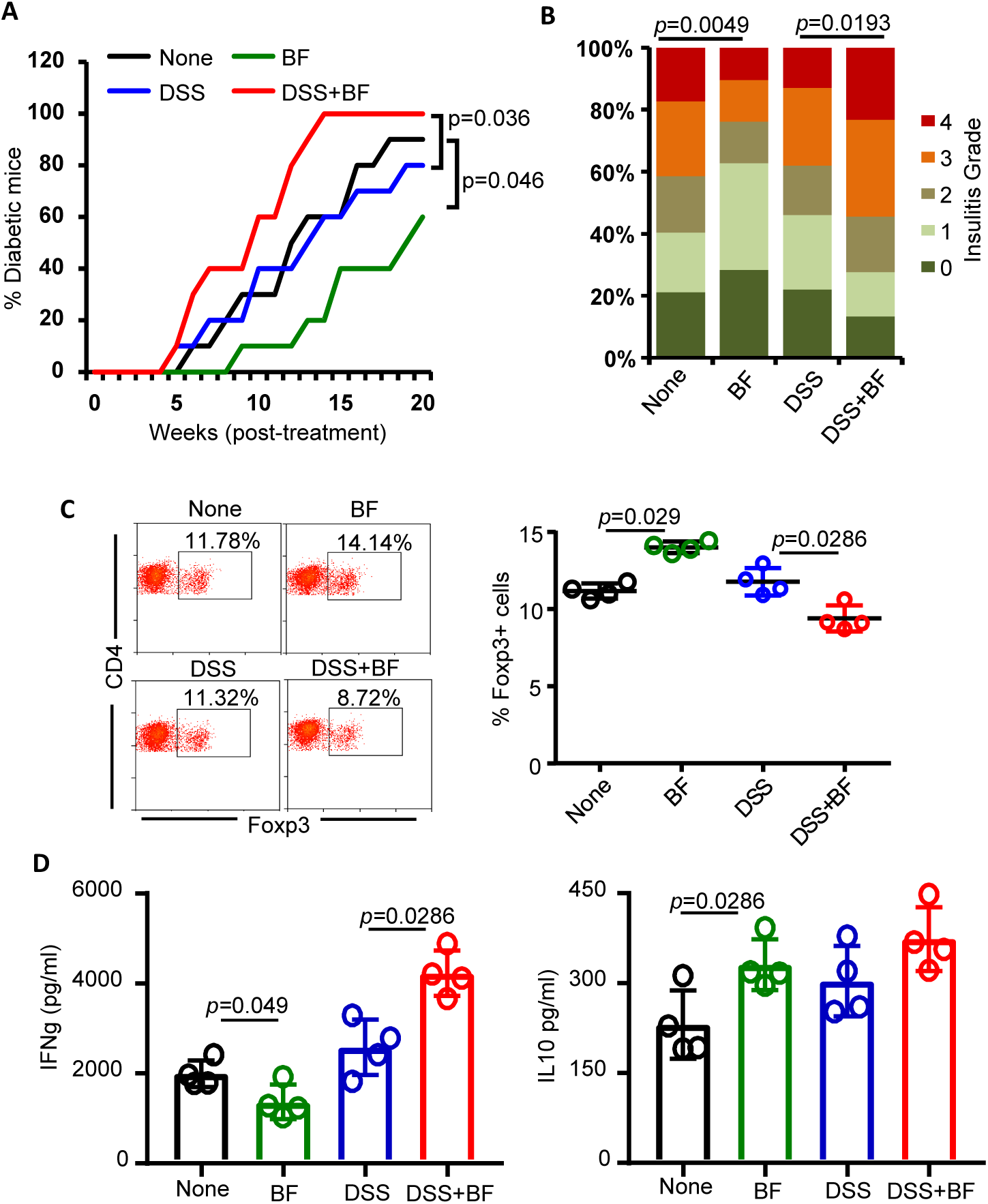
Oral administration of BF under enhanced gut permeability results in accelerated autoimmune progression and early onset of hyperglycemia in NOD mice. Eight week old female NOD mice were maintained on regular drinking water or drinking water containing DSS (0.5%W/V) for 5 days and switched to regular water. A) Cohorts of these mice (n=10/group) were left untreated, or given daily oral gavage of HK BF (500 μg/day/mouse) for 15 days (during and post-DSS treatment), and monitored for hyperglycemia as described for Figs. 1&2. Statistical significance was calculated by log-rank test. **B)** Cohorts of mice (n= 4/group) were euthanized 30 days post-treatment, H&E stained pancreatic tissue sections were examined for insulitis as described for Fig. 1, and the statistical significance was calculated by Fisher’s exact test. **C)** PnLN cells of mice (n=4/group) euthanized 30 days post-treatment initiation were examined for Foxp3+CD4+ T cell frequencies by FACS, and the statistical significance was assessed by Mann-Whitney test. **D)** PnLN cells (n=4 mice/group) were also cultured overnight in the presence of β-cell Ag peptide cocktail for 48 h and the supernatants were tested for cytokine levels by Luminex multiplex assay. The statistical significance was calculated by Mann-Whitney test. Experiment of panels C and D were repeated at least once (using 3 mice/group) with similar statistical trends in results. **Notes:** Fig. S8 shows the degree of gut permeability in low-dose DSS treated and control mice. High dose (2.5% W/V), but not low dose (0.5%), DSS treated NOD mice showed transient loss of body weight, and the weight loss was very modest compared to B6 mice (not shown).

## Discussion

Many years of studies have demonstrated unique immune-modulatory/-regulatory properties and symbiotic nature of a key human colonic bacterium belonging to Bacteroidetes phylum, BF^11, 12^. However, BF is also the primary microbe detected in most clinical intra-abdominal infections/abscesses^15, 16, 36, 44^. This suggested to us that, although the association between intra-abdominal infections and T1D has never been investigated/reported, BF-like human gut commensals may promote immune regulation when restricted to the gut; while their escape to the systemic compartment could produce strong pro-inflammatory response and may impact autoimmune progression. Here, we tested this notion, in a NOD mouse model of T1D, using HK BF and employing oral and i.v. administrations to mimic gut mucosa and systemic exposures to this organism. We also employed a chemical approach to establish enhanced gut permeability and test the translocation of orally administered BF components to the systemic compartment for assessing the potential impacts of systemic exposure to BF on autoimmunity. We show that, in contrast to exposure of gut mucosa to BF which produced protection from T1D, exposure of systemic compartment to BF resulted in a pro-inflammatory response, rapid insulitis progression and early onset of hyperglycemia. Using a PSA deficient mutant of BF and NOD-TLR2-KO mice, we also show that these opposing effects of BF on T1D, upon oral and systemic administrations, are primarily bacterial PSA and host TLR2 dependent.

It has now been established that the gut microbiota is an important environmental factor that modulates T1D disease outcomes^1, 3, 6^. Studies in human subjects and mouse models that have shown the occurrence of dysbiosis very close to disease onset^24, 32^. In general, higher abundance of Bacteroidetes phyla members and a reduction in firmicutes are associated with T1D^23, 24, 28^. However, protection from T1D has also been positively correlated with higher abundance of multiple species belonging to Bacteroidetes^24, 26^. The fact that BF, which is known to colonize the colonic crypt and promote gut immune regulation^11, 18^, is the most commonly isolated gut bacterium in clinical intra-abdominal infections suggests its potential impacts on autoimmune outcomes, especially upon reaching systemic compartment. In fact, higher gut permeability and microbial translocation have been detected in T1D-prone individuals long before the disease onset^41, 45^. Further, it has been shown that mucosal-associated invariant T (MAIT) cells play a critical role in maintaining gut integrity and alterations in the function of these cells, including enhanced cytotoxic properties, which leads to higher gut permeability occurrence in patients with T1D and NOD mice before the onset of clinical disease^46^, supporting the notion that compromised gut integrity and microbial translocation contributes to T1D. It is possible that BF is one of the major gut commensals which gains access to the systemic compartment under enhanced gut permeability in T1D susceptible subjects. However, it is not known if and how the systemic immune response to this gut symbiont contributes to autoimmunity. In this regard, our observations from studies using HK BF in NOD mice, for the first time, show that exposure of the systemic compartment to this agent leads to a profound pro-inflammatory response, diminished Treg function and accelerated autoimmune progression. Our observations from studies using HK BF also substantiate some of the previously reported immune regulatory features^11^ of gut colonization by live BF.

The ability of gut and systemic immune cells to produce different types of immune responses has been studied in the past and the gut mucosa is generally considered tolerant to commensals and dietary antigens^47, 48^. Hence, although the impacts on T1D disease outcomes are intriguing, our observation that distinct immune responses are induced against HK BF by mucosal and systemic immune compartments is not surprising. Nevertheless, it is important to note that not all gut microbial factors promote T1D upon exposing the systemic compartment to them. This notion has been supported by our observation that enhanced gut permeability and translocation of components of normal gut microbiota of NOD mice in our SPF facility alone does not impact T1D incidence. Further, it has been shown that injecting pathogenic bacteria and their components such as lipopolysaccharide (LPS) (TLR4 ligand) induces protection from T1D in NOD mice^49^. Modest impact of systemically administered HK LM and no effect of mutant BF on T1D in NOD mice suggest that our observation on accelerated autoimmune process upon systemic administration is, perhaps, unique to BF-like microbes, and the degree and specificity host-microbe molecular interactions. In this regard, although HK LM, at a dose comparable to that of BF, produced only a modest modulation of T1D in NOD mice, more profound impact upon treatments with higher doses of this microbe cannot be ruled out.

Importantly, the opposing effects of systemic and oral administrations of HK BF in the mouse model of T1D appear to be primarily PSA driven, and its likely interaction with TLR2. BF PSA interacts with TLR2 on APCs as well as T cells and is known to promote immune regulation^11, 19-21^. The ability of PSA to activate conventional and plasmacytoid DCs that further activate/induce Tregs has been described^20^. PSA is also known to be critical for gut colonization by BF and to suppress the colonization by pathogenic bacteria^11^,^30^. Interestingly, it has been shown that not only does TLR2 deficiency protect NOD mice from T1D, TLR2-gut microbiota interaction promotes autoimmunity progression in NOD mice^3, 5, 39^. Hence, our observation that opposing effects on T1D disease outcomes upon exposure of the gut mucosa and the systemic compartment to BF is detectable only in WT, but not TLR2-KO, NOD mice validates the previous reports on the involvement of TLR2 in T1D. Our observations also suggest a paradoxical role for TLR2 signaling in the gut mucosa and in the systemic compartment.

Unlike human gut Gram-negative bacteria such as *E. coli* that produce highly immunogenic LPS, which activates TLR4 signaling, LPS of BF is considered a weak immune activator and does not require TLR4^50^. In fact, we observed that low-dose HK BF produced only modest pro-inflammatory immune response in the systemic compartment as compared to *E.coli* LPS (Fig. S9). Recently, it has been shown that variation in microbiome and LPS immunogenicity can impact T1D in humans^4^. Immune response to *E. coli* LPS, but not the LPS of *Bacteroides* Spp, *B. dorei* particularly, is associated with early immune education and protection from T1D in young children^4^. In agreement with these reports, not only that TLR4 KO NOD mice show rapid onset of hyperglycemia^3^, but also TLR4 binding LPS preparations from Gram-negative bacteria protects NOD mice from T1D^49^. Our observations that oral or systemic administrations of PSA deficient HK BF failed to impact insulitis or hyperglycemia in NOD mice indicate, in addition to the previously reported low immunogenicity and the passive role of its BF LPS, PSA plays a role in modulating the T1D disease outcomes.

Overall, our observations from this study suggest that certain gut microbes such as BF, while promoting immune regulation when restricted to its natural habitat in the distal colon, can contribute to initiation and/or progression of autoimmunity in at-risk subjects upon accessing the systemic compartment. This is evident from the fact that while mice that received HK BF orally under normal conditions were protected from T1D, administration of this agent under enhanced gut permeability accelerated the disease progression. While NOD mice housed in an SPF facility may not encounter many gut permeability inducing events, leaky-gut inducing conditions such as stress, medications and/or infection alone may be sufficient to cause transient microbial translocation and produce inflammatory effects and instigate autoimmunity in at-risk subjects. Importantly, our observations that enhanced gut permeability and microbial translocation alone may not impact the disease outcomes in NOD mice also indicate that specific molecular interactions and immune responses may be critical to influence disease initiation and progression. Of note, higher abundance of Bacteroidetes phyla members including *Bacteroides* Spp has been detected not only in the gut of rodent models of T1D, but also in T1D patients and at-risk children who have progressed towards developing disease symptoms^4, 22-32^, prompting the suggestion that specific *Bacteroides* members could exert pro-autoimmune effects under T1D susceptibility. However, while additional systematic studies in SPF and germ-free NOD mice at different age and disease stages using live BF and other *Bacteroides* members are needed to fully understand, our results obtained using HK BF model and previous studies suggest that colonic symbiotic bacteria such as BF play a critical role in gut and systemic immune regulation under normal circumstances, but could promote autoimmunity under the circumstances such as abnormal epithelial barrier function and higher gut permeability.

## Author contribution

M.H.S. researched and analyzed data, R.G. researched and analyzed data and edited the manuscript, B.M.J researched data, A. J. assisted in experiments, M.C.G. assisted in experiments, and C.V. designed experiments, researched and analyzed data, and wrote/edited manuscript.

## Footnote

This work was supported by unrestricted research funds from MUSC and National Institutes of Health (NIH) grant R21AI133798. Dr. Vasu is the guarantor of this work and, as such, has full access to all the data in the study and takes responsibility for the integrity of the data and accuracy of the data analysis. The authors are thankful to Cell and Molecular Imaging, Pathology, Proteomics, immune monitoring and discovery, and flow cytometry cores of MUSC for the histology service, microscopy, FACS and multiplex assay instrumentation support.

## Data and Resource Availability

The datasets generated during and/or analyzed during the current study are available from the corresponding author on reasonable request.

## Resource availability

No new resources were generated or analyzed during the current study

## Abbreviations

TID: Type 1 Diabetes
NOD: non-obese diabetic
TLR: Toll-like receptor
PSA: polysaccharide A
BF: *Bacteroides fragilis*
ΔPSA: PSA deficient
HK: heat-killed
MLN: mesenteric lymph node
PnLN: pancreatic lymph node
SiLP: small intestinal lamina propria
LiLP: large intestinal lamina propria
PP: Peyer’s patch
DSS: Dextran sulfate sodium

