## Supplemental data for "Polysaccharide A-dependent opposing effects of mucosal and systemic exposures to human gut commensal *Bacteroides fragilis* in type 1 diabetes"

### Fig. S1

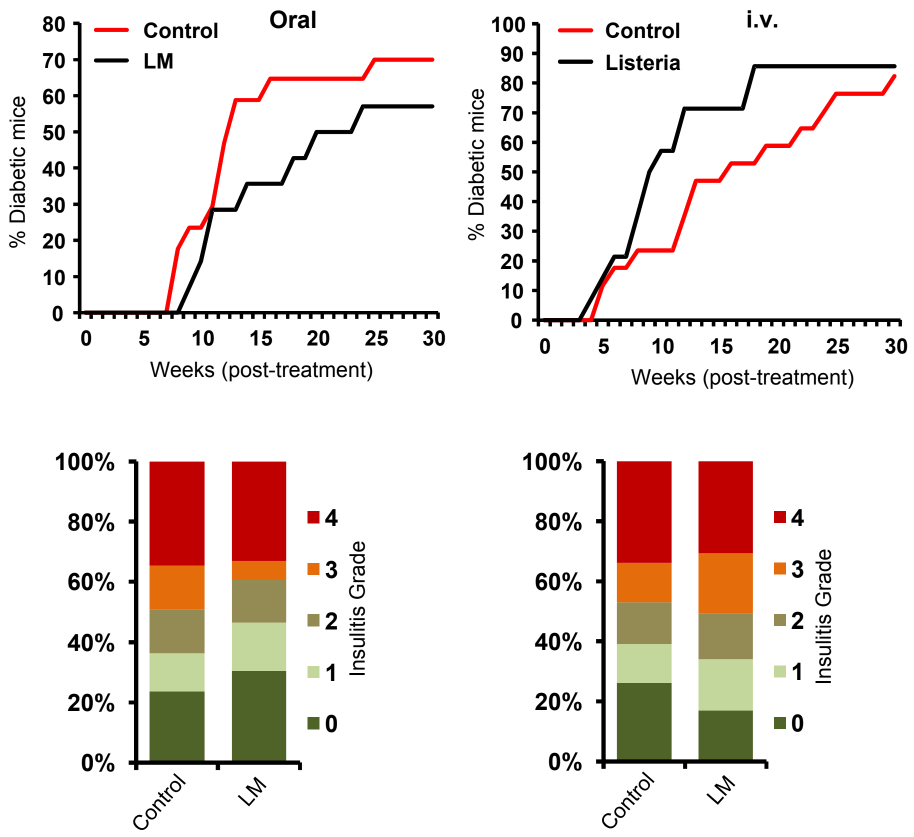


**Fig. S1: Oral or systemic administration of LM resulted in modest modulation of disease progression in NOD mice.** Ten-week-old female NOD mice were given HK LM by oral gavage or by i.v. injection and studied in a similar manner as described for Figs. 1&2 for HK BF. **Upper panels)** Cohorts of control (n=17) and BF treated (n=14) mice were monitored for hyperglycemia as described for Figs. 1 and 2. Statistical significance was assessed by log-rank test. **Lower panels)** Cohorts of mice (n= 5/group) were euthanized 30 days post-treatment and H&E stained pancreatic tissue sections were examined for insulitis as described for Figs. 1 & 2, and the statistical significance was assessed by Fisher's exact test. **Note:** These experiments were carried out in parallel to the experiments of Figs. 1, 2 and 5.

**Fig. S2**

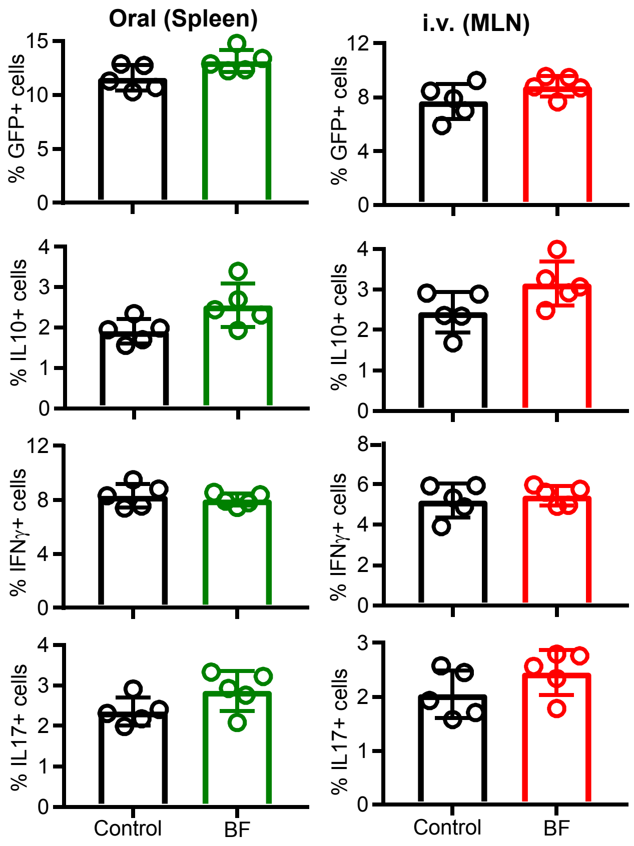


**Fig. S2: Impact of orally and systemically administered BF on T cells of spleen and MLN respectively.** MLN cells of BF injected (systemic) and spleen cells of BF fed (oral), and control NOD-Foxp3-GFP mice described under Fig. 3B (treated with BF for 3 days and euthanized on day 7) were examined for GFP+CD4+ cells, or ex vivo stimulated with PMA and ionomycin for 4 h and examined for intracellular cytokines in CD4+ cells, by FACS. Percentage of cells that are positive for specific marker is shown. n= 5 mice/group and the assay was performed in duplicate for each mouse. This experiment was repeated at least twice (using 3 or 4 mice/group) with similar statistical trends in outcomes. The statistical significance was assessed by unpaired t-test for all panels.

**Fig. S3**

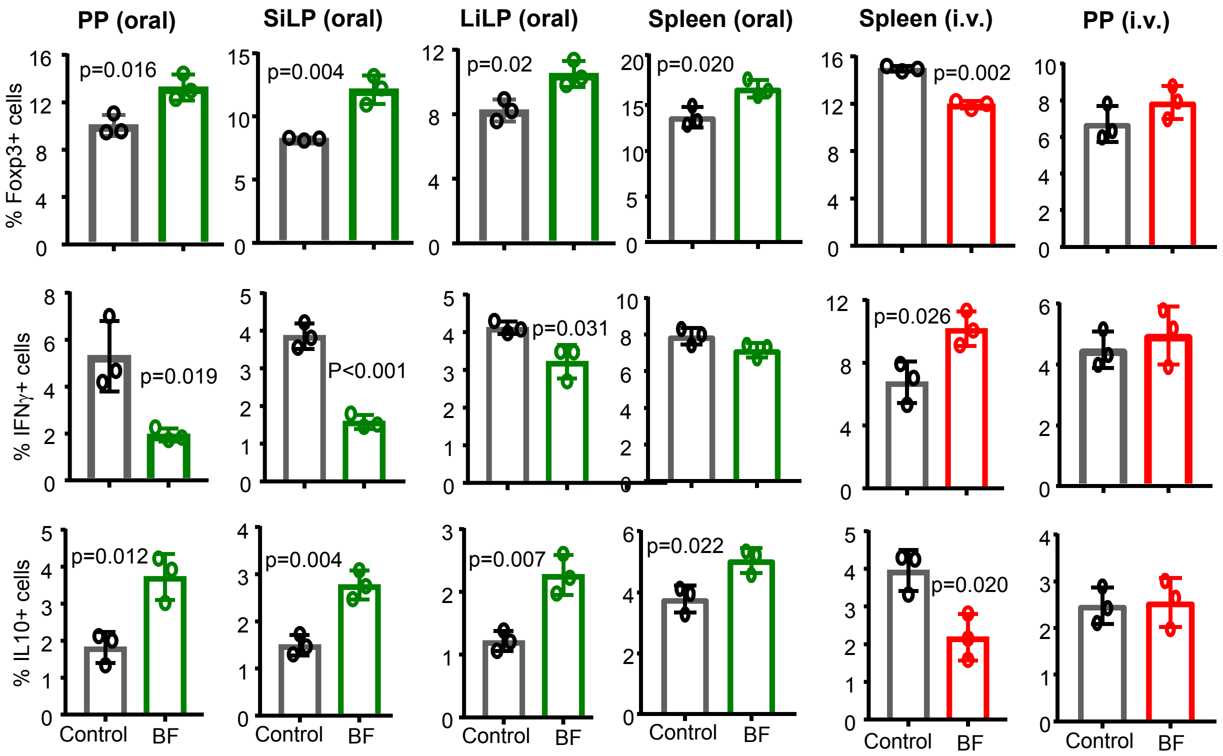


**Fig. S3: Responses of BF exposed immune cells of intestinal and systemic compartments.** Eight week old female NOD mice were given HK BF for 15 consecutive days by oral gavage or every 3 days during a 15 day period by i.v. injection as described for Figs 1 and 2. On day 16, treated and control mice were euthanized, PP, SiLP, LiLP, and spleen cells from mice that received oral treatment and spleen and PP cells from mice that received i.v. injection were examined for Foxp3+, IFNγ+ and IL10+ positive CD4 T cell frequencies by FACS. n= 3 mice/group and the assay was performed in duplicate for each mouse. The statistical significance was assessed by t-test (unpaired; parametric; two-tailed) for all panels. This experiment was repeated once using 3 mice/group with similar statistical trends in results.

**Fig. S4**

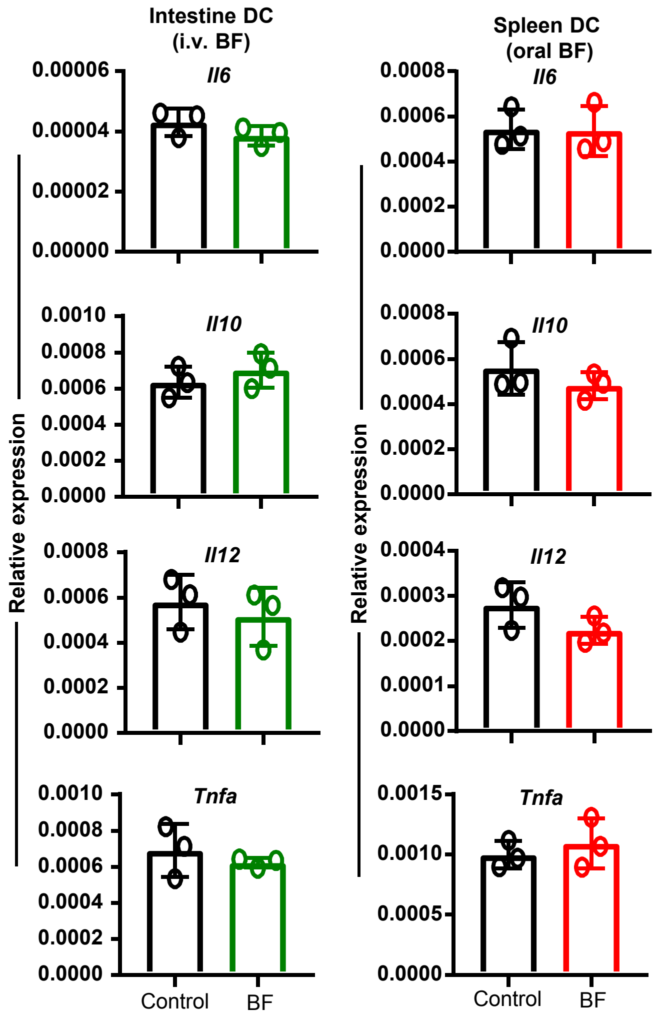


**Fig. S4: Impacts of systemic and oral treatments with BF on intestinal and splenic DCs respectively.** CD11c+ DCs were enriched from the ileum of control and BF injected (systemic) mice and spleen cells of control and BF fed (oral) mice used for Fig. 4A, and subjected to qPCR assay to detect the expression levels of cytokines. n= 3 parallel independent assays using DCs enriched and pooled from 2 mice/group, and each assay was performed in triplicate. The statistical significance was assessed by t-test (unpaired; parametric; two-tailed) . This experiment was repeated once with similar statistical trends in outcomes.

**Fig. S5**

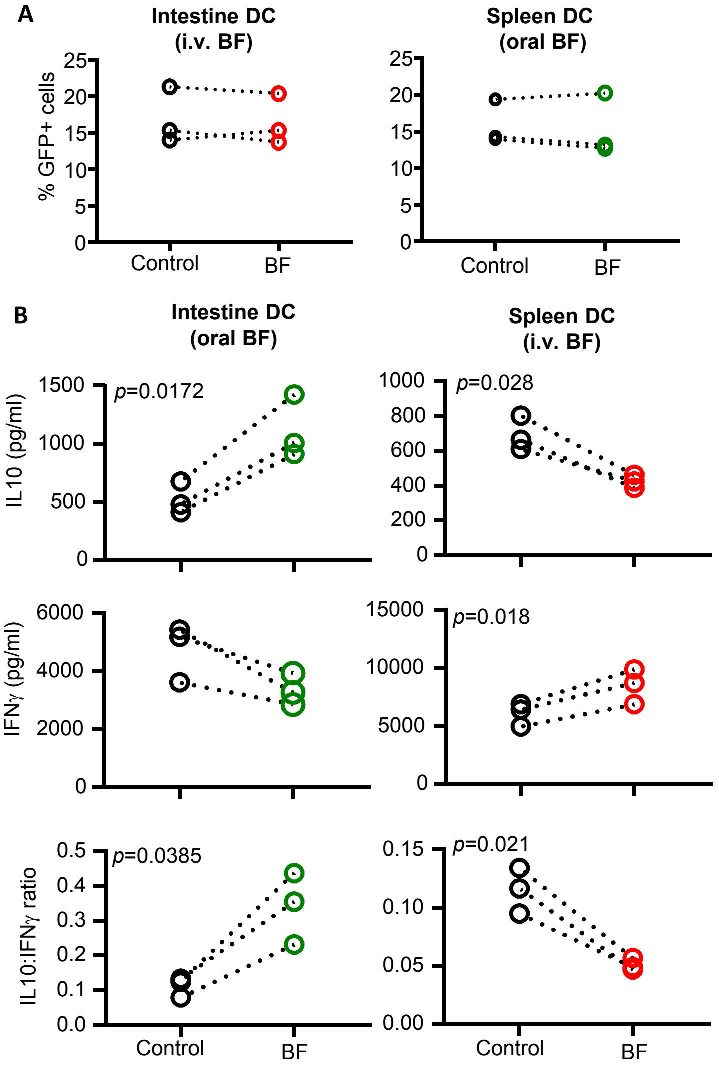


**Fig. S5: Intestinal and splenic DCs from BF treated mice induce different Treg responses and cytokine profiles upon antigen presentation.** Eight week old female NOD mice were given HK BF for 3 consecutive days by oral gavage or by i.v. injection as described for Fig. 3. On day 4, mice were euthanized, CD11c+ DCs that were enriched from ileum and spleen of control and BF fed (oral) mice and spleen and ileum cells of control and BF injected (systemic) mice were pulsed with BDC2.5 peptide and cultured along with T cells from BDC2.5-Foxp3-GFP mice for 4 days as described in Fig. 4B, and examined for GFP+CD4+ cells by FACS. Results from cultures where spleen DCs from BF fed mice and intestinal DCs from BF injected mice are shown in **panel A**. Results from cultures where intestinal DCs from BF fed mice and spleen DCs from BF injected mice are shown in Fig. 4B. Supernatants from selected cultures were tested for cytokine levels by ELISA **(panel B)**. n=3 independent experiments using DCs enriched and pooled from 2 mice/group in each experiment. Each assay was performed in triplicate, values were averaged, and the cumulative values of three assays are shown. The statistical significance value was determined by t-test (paired; parametric; two-tailed) which compared control and BF groups within each assay as indicated by the dotted lines. Supernatants from cultures of spleen DCs from BF fed mice and intestinal DCs from BF injected mice showed no significant differences in the cytokine profiles compared to respective control DCs (not shown).

### Fig. S6

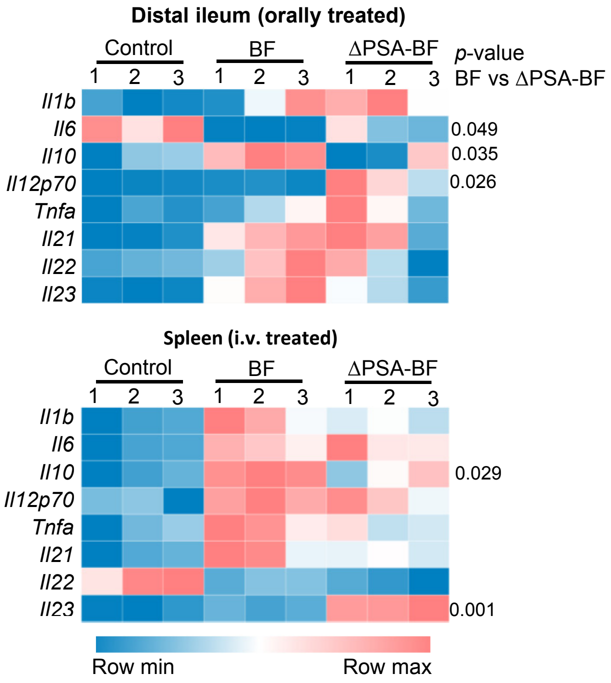


**Fig. S6: PSA deficiency of BF impacts cytokine responses by the gut mucosa and systemic immune cells.** Eight week old female NOD mice were given HK BF or ΔPSA-BF for 3 consecutive days by oral gavage (500 µg/mouse/day) or by i.v. injection (10 µg/mouse/injection). On day 4, mice were euthanized and cDNA prepared from distal ileum of control and BF fed mice and spleen cells from control and BF injected mice were subjected to qPCR assay and the expression levels of cytokines were compared. Expression levels relative to β-actin expression were plotted as heatmaps. n= 3 mice/group and the assay was performed in triplicate for each mouse. BF and ΔPSA-BF groups were compared by t-test (unpaired; parametric; two-tailed) for assessing the statistical significance. This experiment was repeated using 3 mice/group at least once with similar statistical trends in outcomes.

### Fig. S7

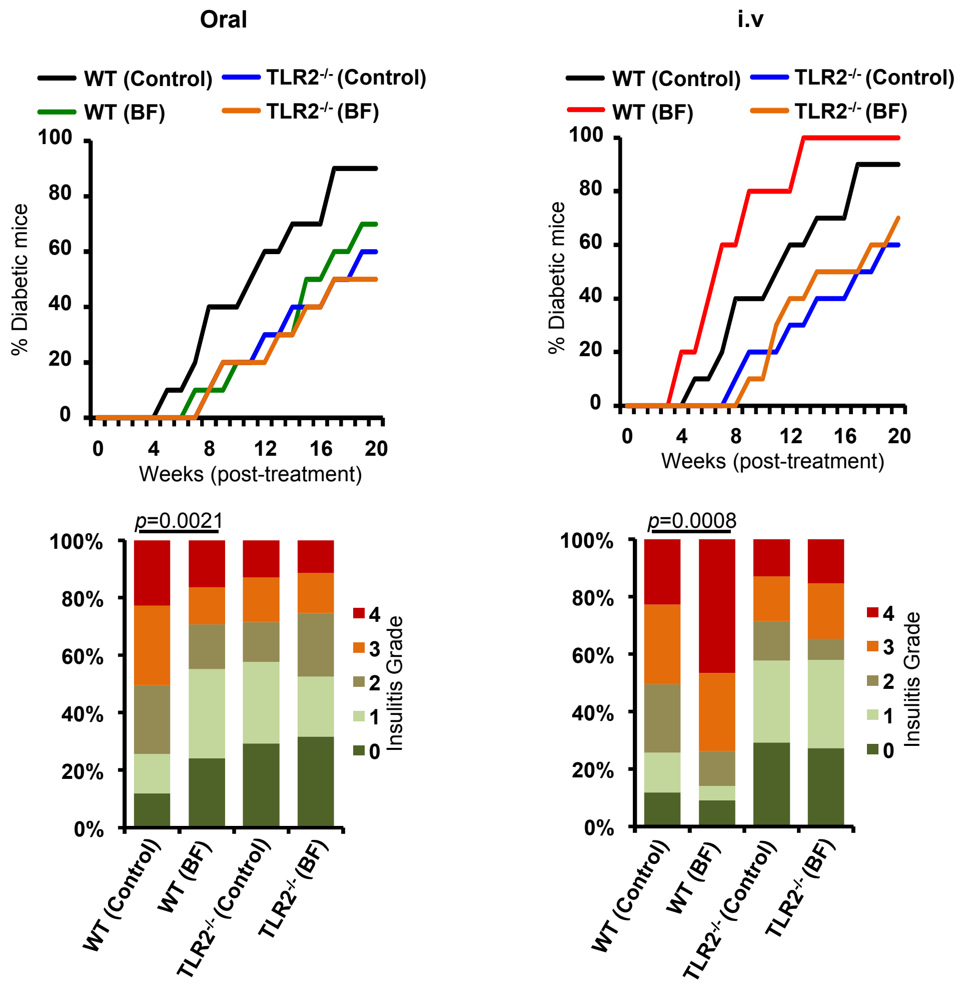


**Fig. S7: Opposing T1D modulatory effects of oral and systemic administration of BF were observed only in WT, but not TLR2 KO, NOD mice.** Ten to twelve-week-old female WT-NOD and TLR2$^{-/-}$ NOD mice were given HK BF by oral gavage or by i.v. injection and studied as described for Figs. 1&2. **Upper panels)** Cohorts of control (n=10) and BF treated (n=10) mice were monitored for hyperglycemia as described for Figs. 1 and 2. Control and treated groups of WT and TLR2$^{-/-}$ mice were compared and the statistical significance was determined by log-rank test. **Lower panels)** Cohorts of mice (n= 3/group) were euthanized 30 days post-treatment and H&E stained pancreatic tissue sections were examined for insulitis as described for Figs. 1 & 2, and the statistical significance was assessed by Fisher's exact test.

**Fig. S8**

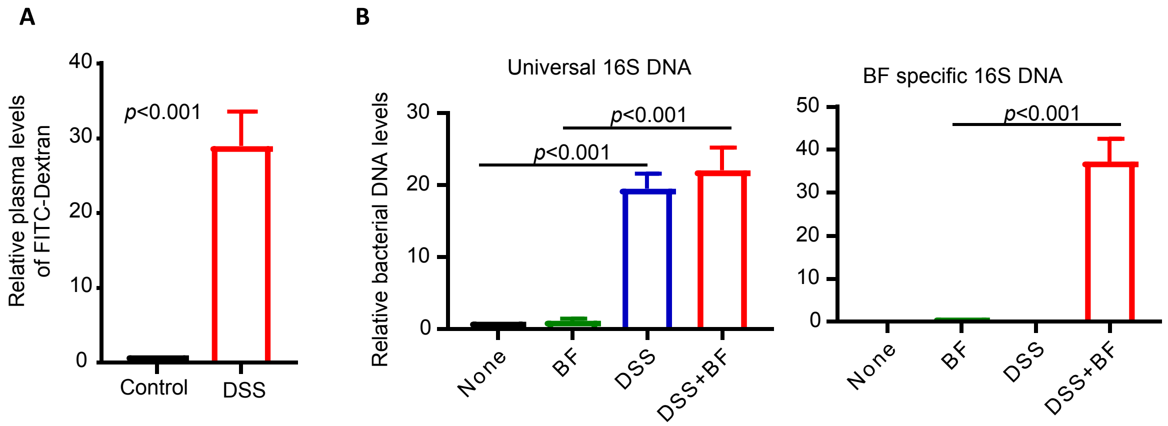


**Fig. S8: Enhanced gut permeability in DSS treated mice. A)** Pre-diabetic NOD mice were left alone or treated with DSS in drinking water (0.05% W/V) for 5 days and switched to regular water. These mice (n=3/group) were given FITC-Dextran (10 µg/mouse) by oral gavage on day 6, euthanized after 4 h, and the plasma samples were examined for FITC-Dextran concentration using a fluorescence microplate reader against control plasma samples spiked with known concentration of FITC-Dextran. Mean value from the untreated group was considered as 1 and the fold FITC-Dextran levels in DSS treated mice were calculated against this baseline. **B)** Pre-diabetic NOD mice were given DSS in drinking water for 5 days, switched to drinking water for 24 h. Cohorts of mice received HK BF (500 µg/day/mouse) by oral gavage all 6 days. These mice (n=3/group) were euthanized 4 hours post-treatment on day 6, blood samples were tested for total bacterial DNA (using universal 16S rDNA primers) and BF specific DNA (using BF 16S rDNA primer sets) by qPCR. For left panel, relative total bacterial DNA levels were calculated against the none group and compared with the respective control group. For right panel, BF specific DNA levels were calculated and the values of BF and DSS+BF groups were compared. The statistical significance value was determined by t-test (paired; parametric; two-tailed) . These experiment were repeated once (using 3 mice/group) with similar statistical trends in results.

### Fig. S9

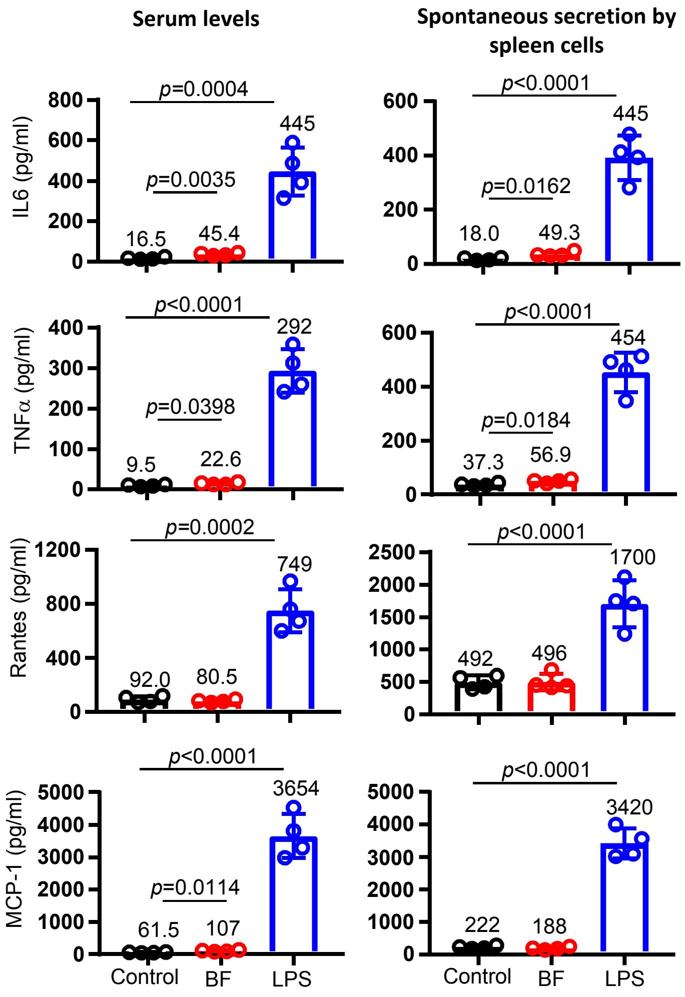


**Fig. S9: Systemic treatments with HK BF, compared to Bacterial LPS, produces only a mild pro-inflammatory response.** Female NOD mice (8-10 week old; 4 mice/group) were injected i.v. with HK BF (10 µg/mouse/day) or *E.coli* LPS (1 µg/mouse/day) for 3 consecutive days and euthanized on day 4 to obtain serum and spleen cells. Spleen cells ($2x10^6$ cells/ml) were cultured overnight without additional stimulus and the spent media were collected. Pro-inflammatory cytokine and chemokine levels in serum and culture media were examined by Luminex multiplex assay and the concentration of selected factors are shown. The statistical significance was assessed by t-test (unpaired; parametric; two-tailed) .
